# Multi-grip classification-based prosthesis control with two EMG-IMU sensors

**DOI:** 10.1101/579367

**Authors:** Agamemnon Krasoulis, Sethu Vijayakumar, Kianoush Nazarpour

## Abstract

In the field of upper-limb myoelectric prosthesis control, the use of statistical and machine learning methods has been long proposed as a means of enabling intuitive grip selection and actuation. Recently, this paradigm has found its way toward commercial adoption. Machine learning-based prosthesis control typically relies on the use of a large number of electrodes. Here, we propose an end-to-end strategy for multi-grip, classification-based prosthesis control using only two sensors, comprising electromyography (EMG) electrodes and inertial measurement units (IMUs). We emphasize the importance of accurately estimating posterior class probabilities and rejecting predictions made with low confidence, so as to minimize the rate of unintended prosthesis activations. To that end, we propose a confidence-based error rejection strategy using grip-specific thresholds. We evaluate the efficacy of the proposed system with real-time pick and place experiments using a commercial multi-articulated prosthetic hand and involving 12 able-bodied and two transradial (i.e., below-elbow) amputee participants. Results promise the potential for deploying intuitive, classification-based multi-grip control in existing upper-limb prosthetic systems subject to small modifications.

## I. INTRODUCTION

Upper-limb myoelectric prostheses are electromechanical devices that aim to partially restore the functionality and appearance of a missing limb. They typically comprise a muscular activity recording unit based on surface electromyography (EMG), an active end-effector, such as a prosthetic hand with motorized digits and a wrist rotation unit, and a processing unit that translates the recorded muscular activity information into motor commands for the end-effector. Nowadays, there exists a plethora of prosthetic devices with remarkable mechanical properties, nearly approximating the dexterity of the human hand. Nevertheless, the full mechanical capabilities of state-of-the-art prosthetic hands are seldom utilized; the ineffectiveness of the deployed control algorithms, the non-stationary nature of EMG signals, and lack of an ample number of independent muscle control sites hinder the control dexterity of myoelectric prostheses [1].

State-of-the-art active prostheses, such as the Össur i-Limb ultra [2] and Ottobock bebionic [3] hands, are typically shipped with a set of pre-programmed grip patterns. These can be utilized by the user to hold objects or perform other activities of daily living. A pair of surface EMG electrodes is commonly used to monitor the activity of flexor and extensor muscle groups, thus providing the user with control over a single degree of freedom (DOF), such as hand opening/closing within a specified grip configuration or wrist rotation. To switch between different grip patterns or functions, the user has to either perform a series of muscle co-contractions to shift through the available modules [4] or send a trigger signal associated with the desired grip, such as a double or triple impulse. This control scheme is robust but lacks intuitiveness, which in turn can lead to increased cognitive load.

Since the 1970’s, significant efforts have been put into utilizing computational tools from the fields of statistics and machine learning to enhance the control of upper-limb myoelectric prostheses. One prominent example of this approach is the use of classification algorithms with the aim of increasing the intuitiveness of the control interface. The fundamental working principle of this paradigm is that features extracted from multiple EMG electrodes form motion-specific clusters in a high-dimensional space that can be used to discriminate classes of movement. Thus, to access a specific function, a user only needs to activate their muscles in a naturalistic fashion, much like they would do with an intact limb. This approach has demonstrated proof-of-principle for decoding arm movements, such as elbow and wrist flexion/extension and hand opening/closing [5], [6], and has recently found its way to commercial adoption [3], [7]. Several studies have also used this paradigm to decode grips and gestures for intuitive prosthetic hand control. In their majority, however, they have been either limited to offline analyses [8]–[10] or only included able-bodied participants [11]–[13], with few exceptions demonstrating real-time control with amputees [14]–[16].

Farina *et al.* [1] have identified the ability to achieve high decoding performance with a minimal number of electrodes as one of the main challenges for machine learning-based myoelectric control. A significant body of work has previously investigated means of achieving this goal. Exhaustive search or sequential selection algorithms have been used to identify a suitable subset from a larger pool of sensors, typically in the range of 4 to 12 [14], [17]–[25]. Despite previous efforts, the feasibility of using a single pair of sensors, which is typically available in commercial prostheses, to control a multi-grip prosthetic hand has yet to be demonstrated.

Drastically reducing the amount of sensors used for myoelectric control may lead to a decrease in classification performance. Additionally, it has been reported that unintended prosthesis motions can lead to user frustration [26], which in turn may increase the risk of prosthesis rejection. Thus, to ensure user satisfaction, it is imperative to design fault-tolerant myoelectric controllers with the ability to reject classification predictions unless they are made with high confidence. Several post-processing strategies have been proposed for reducing the frequency of unintended prosthesis activations, including but not limited to: majority voting, whereby the control action at a given time step is affected by previous and potentially future predictions (i.e. non-causal filter) [27]; training multiple binary one-vs.-rest or one-vs.-one classifiers and rejecting classification predictions unless unanimous agreement is reached among the pool of classifiers [26], [28]; threshold-based confidence rejection, whereby a classification prediction is rejected unless the respective posterior probability exceeds a fixed [6], [14], [29] or class-specific [24] threshold; and using an auxiliary binary classifier to estimate whether the predictions of the base motion classifier are accurate [30]. All these methods reduce the rate of unintended prosthesis activations. However, this usually happens at the expense of an increase in computational complexity [26], [28], [30], response delay [27] or a decrease in overall classification accuracy [26].

In this work, we propose an end-to-end pipeline for realtime prosthetic hand control for transradial (i.e., below-elbow) amputees by using two sensors comprising EMG electrodes and inertial measurement units (IMUs). Special attention is given to optimizing system parameters such that the amount of unintended performed motions is minimized. To that end, a novel algorithm is introduced for selecting class-specific confidence thresholds based on false positive rate minimization. The efficacy of the system is evaluated with object pick and place experiments using a commercial prosthetic hand and involving both able-bodied and transradial amputee participants. To our knowledge, this is the first time that machine learning-based, multi-grip prosthesis control is demonstrated in realtime with amputees by using only two EMG-IMU sensors.

## II. METHODS

### A. Participant recruitment

Twelve able-bodied (10 male, two female; 10 right-hand, two left-hand dominant; median age 28 years) and two right-arm, transradial amputee subjects were recruited. Both amputee participants were right-hand dominant prior to amputation. Some of the able-bodied and both amputee subjects had previously taken part in classification-based myoelectric control experiments [14]. Experimental procedures were in accordance with the Declaration of Helsinki and were approved by the local Ethics Committees of the School of Informatics, University of Edinburgh (#201507160854) and School of Engineering, Newcastle University (#14-NAZ-056). Participants read an information sheet and gave written informed consent.

### B. Signal acquisition and socket fitting

For the able-bodied group, 16 EMG-IMU Delsys^®^ Trigno™ IM sensors were placed on the participants’ forearm arranged in two rows of eight equally spaced sensors each (Fig. 1). For the two amputee participants, 13 and 12 sensors were used, respectively, due to limited space availability. The sensors were placed on the able-bodied participants’ dominant arm, whereas for amputees they were placed on the subjects’ stump (right arm in both cases). Prior to sensor placement, the participants’ skin was cleansed using 70% isopropyl alcohol. Elastic bandage was used to secure the sensor positions throughout the experimental sessions. Following sensor placement, we verified the quality of all EMG channels by visual inspection. The hardware sampling rates for EMG and inertial data were 1111 Hz and 128 Hz, respectively. The IMU components comprised 3-axes accelerometers, gyroscopes and magnetometers, measuring respectively acceleration, angular velocity and orientation. Readings from IMUs were used in their raw format, therefore no calibration was required.

**Fig. 1.**
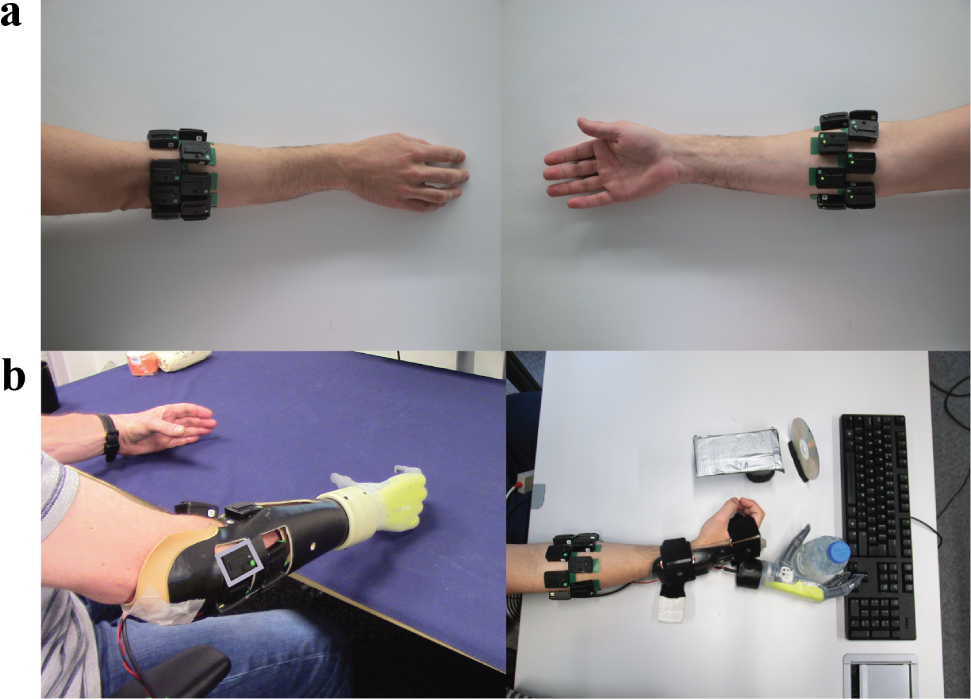
Sensor placement and experimental setup. **a**, Two rows of eight equally-spaced sensors were placed around the forearm of the able-bodied participants. For the two amputee subjects, 12 and 13 sensors were used, respectively. For each participant, the optimal subset of two sensors was identified and used for real-time decoding. **b**, Pictures of one amputee *(left)* and one able-bodied *(right)* participant during the training and real-time control phases of the experiment, respectively. The shown amputee participant performed bilateral mirrored movements during training. For able-bodied participants, a custom-built splint was used to support the prosthesis. The two amputee participants wore modified own sockets. A grey box is drawn around one of the sensors attached to the amputee participant’s arm.

A splint and custom built sockets were designed to accommodate the prosthetic hand that was used in the experiments (Fig. 1). The same splint was used for all able-bodied participants and was adjusted using Velcro straps. The two amputee participants wore their own sockets, which were modified to accommodate the EMG-IMU sensors. The prosthesis was fitted on the dominant arm of the able-bodied subjects. For the two amputees, it was fitted on their stump.

### C. Robotic hand

The Össur robo-limb™ hand was used in the experiments, which is an externally-powered, underactuated (11 DOFs, 6 degree of activations (DOAs)), anthropomorphic hand. This model is identical to the commercially available “i-Limb ultra” model. It comprises 5 motors controlling the flexion/extension of the digits and an additional motor controlling the rotation of the thumb. The hand operates under 7.4 V nominal voltage with a maximum current consumption of 7 A. The hand was externally powered with a doubly-insulated power supply unit, certified for medical experiments.

The robo-limb can be controlled by a computer via a CAN bus interface in an open-loop fashion. The control commands take the form “`ID` - `Action` - `PWM`”: `ID` specifies the desired DOA to be activated (0-6); `Action` indicates the desired motion (open/close/stop); and `PWM` specifies the desired pulse width modulation level to be applied to the motor, in the range [10, 127], which controls the digit movement velocity.

Grip control was implemented for the hand using a set of pre-defined digit activation sequences. A grip was executed only if the most recent grip command had finished execution. In any other case, that is, if a new command was issued while the most recent one was still being executed (i.e. digits were moving), the newly issued command would be ignored.

### D. Behavioural task

The participants sat comfortably on an office chair. Each session comprised a calibration and a real-time control phase. In the *calibration* phase, subjects were instructed to perform five motions presented to them on a computer monitor: power grasp, lateral grasp, tripod grasp, index pointer, and hand opening. For each calibration trial, participants were instructed to execute the respective movement at a moderate speed. Once they had executed the desired grip, they were required to perform a dynamic movement, thus covering with their (residual) arm the region of interest. This approach was followed because it has been previously shown that it can help alleviate the limb position effect [31] and improve decoding performance [14], [15]. One of the amputee participants performed bilateral mirrored movements as they found it was easier in that way to perform phantom limb muscle activations. Calibration trials lasted for 5 s and were interleaved with 3 of rest. Two separate blocks of data were recorded, each one comprising 10 consecutive repetitions for each grip. The two datasets were subsequently used as *training* and *validation* sets, respectively.

In the interval between calibration data collection and realtime control, a subject-specific optimal subset of two EMG-IMU sensors was identified (see Section II-G) and used to train a regularized discriminant analysis (RDA) classifier (see Section II-H). Class-specific confidence thresholds were also estimated at this stage (see Section II-I).

In the *real-time control* phase, participants were instructed to use the prosthetic hand to grasp, relocate (approximately 50 cm away from initial position), and release three objects, and finally press the space bar on a computer keyboard. The following four objects were used: a water bottle, a credit card simulator, a CD, and a computer keyboard. The corresponding prosthesis grips were power, lateral, tripod, and pointer. To switch between different grips, participants first had to fully open the hand. Depending on their laterality, participants were instructed to move the objects away from their point of reference; from center to right for the right-handed able-bodied and the two amputee subjects, and from center to left for the left-handed able-bodied subjects. Participants were instructed to complete the trials as fast as they could and trial timings were recorded by the experimenter. Trials were considered successful if all objects were relocated and the space key was pressed within 75 s. In case of an object drop during relocation, the trial would be interrupted and considered unsuccessful. The number of trials per subject was set to 10 and participants were given 45 s of rest in-between trials. The object presentation order was pseudo-randomized and counter-balanced across participant groups. During real-time control, participants were blind to the number of sensors used for decoding.

### E. Performance assessment

For offline analyses, the multi-class *cross-entropy loss* — also known as *logistic loss* or *log-loss*— was used to evaluate decoding performance. The cross-entropy loss is closely related to the *Kullback-Leibler divergence* between the empirical and estimated distributions of a discrete random variable. Let *y* ∈ 1,…, *C* denote a discrete target variable which is encoded as a “one-of-K” binary indicator matrix *Y* of dimensionality *N* × *C*, such that:

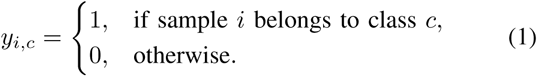

The multi-class cross-entropy loss is then defined as follows:

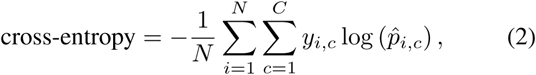

where 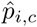 denotes the posterior probability that sample *i* belongs to class *c*. In the ideal case—that is, when all samples in a dataset are correctly classified and the corresponding posterior probabilities are exactly 1— the cross-entropy loss is equal to 0. On the other hand, there is no lower bound for cross-entropy loss; poor probability predictions can yield arbitrarily low (i.e., large negative) scores. This metric was selected for performance evaluation because, as opposed to classification accuracy, it can also assess the quality of posterior probability estimates. Prior to performance evaluation, the distribution of validation/test samples was balanced by undersampling the dominating “rest” class.

To evaluate prosthetic control performance in the real-time experiments, we adopted the following two common task-related metrics: completion rate, that is, the ratio of successful to total number of trials; and completion time, defined as the time taken to accomplish a successful trial.

### F. Signal pre-processing

Power line interference was suppressed from EMG signals by applying a Hampel filter. To remove motion artefacts, band-pass filtering was applied in the range [10, 500] Hz using a 4^th^-order Butterworth filter. Myoelectric and inertial signals were synchronized via upsampling to 2 kHz and linear interpolation.

By using a sliding window approach, we extracted four time-domain EMG features, namely, waveform length, Wilson amplitude, log-variance, and 4^th^-order auto-regressive coefficients; a total of seven attributes per EMG channel. For inertial data (i.e., 3-axis acceleration, angular velocity, and magnetic field), the mean value within the processing window was used; that is, nine attributes per IMU were extracted. The length of the sliding window was set to 128 ms and the increment to 50 ms (60% overlap). For offline training, the total number of extracted features was 256 for able-bodied subjects, 208 for the first, and 192 for the second amputee participant, respectively (i.e., 16, 13, or 12 sensors × 16 features/sensor). The input features were standardized via mean subtraction and inverse standard deviation scaling. For real-time decoding, only two sensors were used for each participant (see Section II-G), therefore the number of features was 32.

Stimulus presentation timings were recorded using high resolution timestamps. We used a post-hoc relabelling procedure based on EMG-stimulus alignment to refine the exact motion timings and target vector labels for each subject and trial [32].

### G. Sensor selection

For each subject, two EMG-IMU sensors were selected from the full set by using a sequential forward selection algorithm. As part of this procedure, we fitted linear discriminant analysis (LDA) models using the training set and assessed performance on the validation set using cross-entropy loss. In each iteration of the search algorithm, a sensor could be added to the selected subset using all 16 relevant features.

### H. Classifier training and hyper-parameter optimization

For movement intent decoding, we used the RDA classifier, which offers a continuum between LDA and quadratic discriminant analysis (QDA) by fitting class-specific covariance matrices that are regularized toward the pooled covariance matrix [33]. To tune the regularization hyper-parameter, we used a line search in the range [0, 1] with a step size of 0.025 and selected the parameter value that yielded the lowest cross-entropy loss on the validation set. Following hyper-parameter optimization, the training and validation sets were merged and used to train final models. Model training and hyper-parameter optimization were performed in a subject-specific fashion.

For the purposes of post-hoc offline analysis, a slightly different model selection and evaluation approach was used. In this case, performance was assessed using 10-fold cross-validation on the second dataset for each participant, by using a 90%-10% split. That is, the whole of the first dataset for each individual was used to train models (*training set*), 9 out of 10 repetitions of each motion from the second dataset were used as a *validation set* for sensor selection and RDA hyper-parameter optimization, and the collection of left-out repetitions from the second dataset were used as a *test set*.

### I. Confidence-based rejection and threshold selection

Classifier predictions were post-processed using confidence-based rejection. That is, predictions were discarded unless the corresponding posterior probabilities exceeded pre-defined, class-specific thresholds. Furthermore, when the “rest” class was predicted by the decoder, there was no movement and the hand held its previous state. The rejection thresholds were selected by using receiver operating charachteristic (ROC) analysis on the validation set. To that end, multiple one-vs.-all RDA classifiers were trained and the corresponding false positive rate and true positive rate scores were computed for threshold values in the range [0, 1]. The rejection threshold for each class was selected such that the true positive rate was maximized, while the respective false positive rate was constrained to be smaller than a cut-off value, set *a priori* to 5 × 10^−4^. This was done to minimize the number of false positives that would translate into unintended hand motions. To avoid setting thresholds extremely close to 1 for well-separated classes, which would dramatically reduce the respective true positive rate in real-time control, we introduced an upper-bound for the thresholds and set it empirically to 0.995.

### J. Statistical analysis

All statistical comparisons were performed using non-parametric tests. Two-sided Wilcoxon signed-rank tests were used for pairwise comparisons and multiple comparison tests were corrected using the Šidák method. The statistical significance level was set to *α* = 0.05.

## III. RESULTS

### A. Offline analysis

We evaluated decoding performance for the family of discriminant analysis classifiers, namely, LDA, RDA, and QDA, with a varying number of sensors. The results of the classifier comparison analysis are presented in Fig. 2a. Performance was assessed using the cross-entropy loss (see Eq. (2)), with lower values indicating better decoding performance. In general, performance improved as new sensors were added to the decoders and reached a plateau after the inclusion of 6-8 sensors. The RDA classifier outperformed LDA for small numbers of sensors, but the two algorithms yielded comparable scores for more than five sensors. The performance of QDA was remarkably worse than that of LDA and RDA. Notably, the performance of QDA deteriorated when a large number of sensors was used. The results were consistent across the able-bodied and amputee populations.

**Fig. 2.**
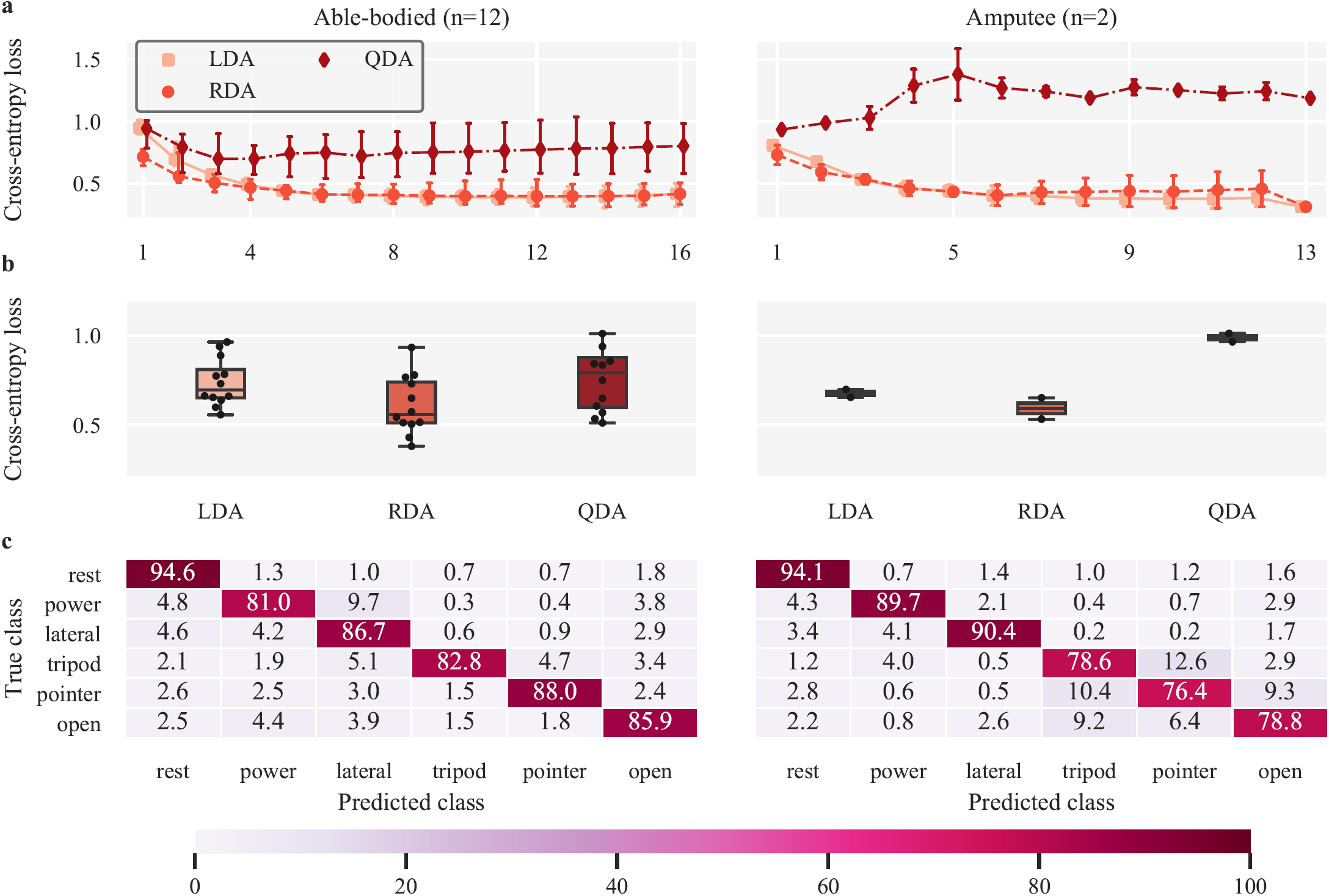
Offline analysis. **a**, Classifier comparison for a varying number of EMG-IMU sensors. The three considered classifiers were LDA, QDA, and RDA. The metric used for comparison was the cross-entropy loss (i.e., log-loss); lower values indicate better performance. For each configuration, the optimal subset of sensors was identified using sequential forward selection. A total of 16 features were extracted from each sensor (seven EMG and nine IMU features). For amputees, 12 sensors were used for the second participant, thus a single data point is shown in traces corresponding to 13 sensors and confidence intervals are not estimated. Points, medians; error bars, 95% confidence intervals estimated via bootstrapping (1000 iterations). **b**, Detailed classifier comparison when using the optimal subset of two sensors. Straight lines, medians; solid boxes, interquartile ranges; whiskers, overall ranges of non-outlier data; dots, individual data points. **c**, Average confusion matrices for the two participant groups using the RDA classifier and the optimal pair of sensors. Annotated scores and color intensities indicate average per-class normalized accuracy scores.

The results from using the optimal subset of two sensors are presented in detail in Fig. 2b, separately for the able-bodied and amputee groups. A Friedman test revealed a statistical effect of decoding algorithm on performance (*p* = 10^−3^, *n* = 14). Post-hoc pairwise comparisons showed that the RDA classifier significantly outperformed LDA and QDA (*p* = 10^−2^ in both cases; *n* = 14), whereas LDA performed marginally, but not significantly, better than QDA (*p* = 0.18).

Average confusion matrices with the two optimally selected EMG-IMU sensors and the RDA classifier are shown for each participant group in Fig. 2c. The annotated scores indicate average per-class normalized accuracy scores for each group.

### B. Real-time myoelectric prosthesis control experiment

The working principle of the proposed real-time prosthetic control paradigm is illustrated in Fig. 3 using a representative trial with one participant. The raw EMG signals from the two sensors selected for this subject are shown in Fig. 3a. The time series of classification prediction and prosthesis activation are shown in Fig. 3b and the temporal evolution of the class posterior probability distribution is shown in Fig. 3c. For the shown trial, the sequence of objects to be relocated was “card”, “bottle”, and “CD”; thus, the required sequence of hand grips was “lateral”, “open’, “power”, “open”, “tripod”, “open”, and “pointer”. It can be observed from Fig. 3b that although there was a relatively large number of incorrectly classified instances (black trace), the confidence-based rejection strategy discarded most of them as the corresponding posterior probabilities did not exceed the respective class-specific confidence thresholds (Fig. 3c). Overall, there were two unintended hand motions in this trial, marked with red ellipses in Fig. 3b, and the trial was successful with a completion time of 26.9 s.

**Fig. 3.**
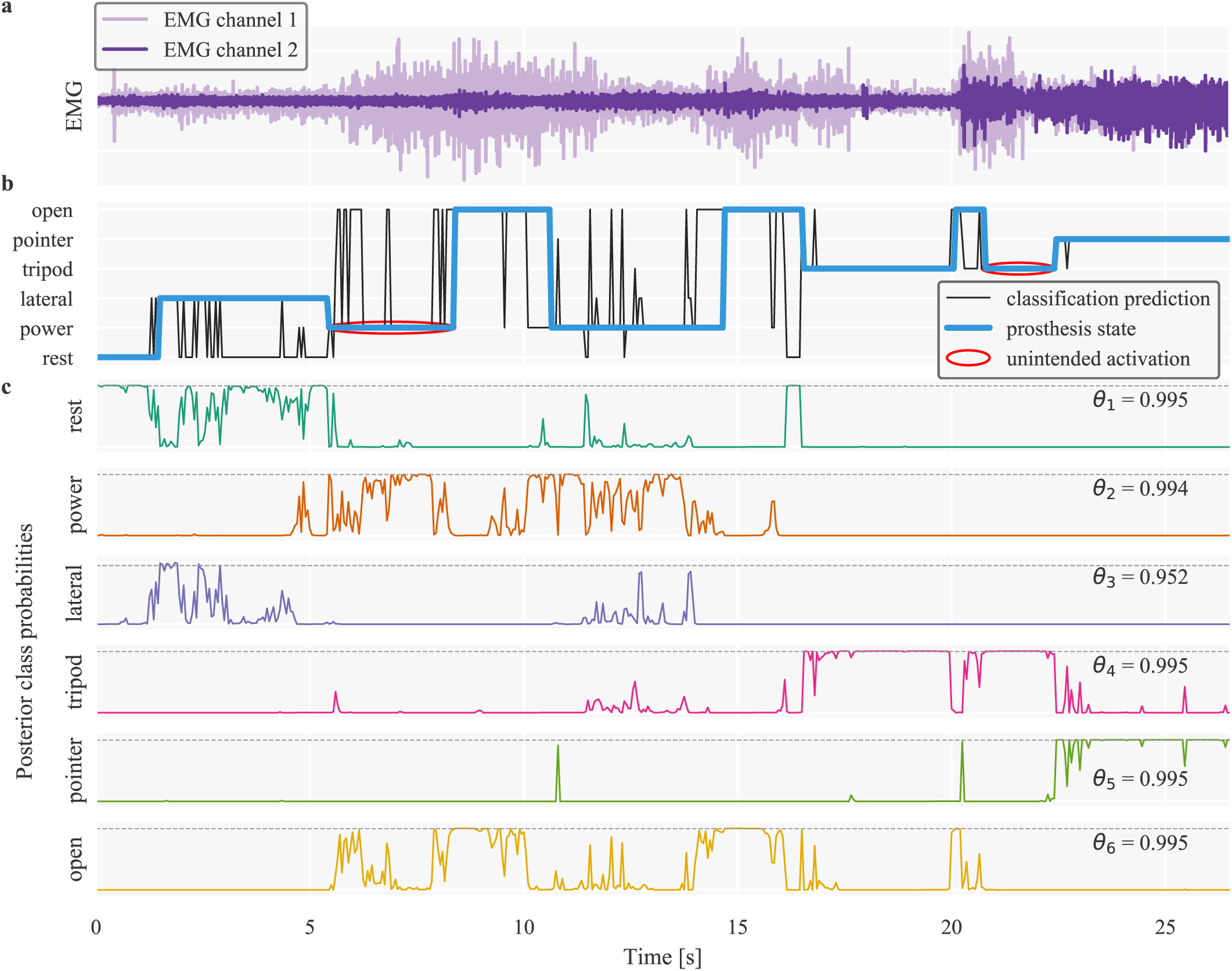
Working principle of the proposed real-time prosthesis control framework. A representative trial is shown for one participant. The object presentation order for this trial was “card”, “bottle’, and “CD”; thus, the desired grip activation sequence—including the required intermediate hand “open” triggers to switch between grasps—was “lateral”, “open”, “power”, “open”, “tripod”, “open”, and “pointer”. **a**, Time-series of raw EMG signals from the two sensors used for the participant. **b**, Time-series of classifier predictions and temporal evolution of prosthesis state. A new classification was translated into a control action only if the corresponding posterior probability exceeded the respective class-specific threshold. When the posterior probability was lower than the corresponding rejection threshold, or the “rest” class was predicted, the hand held its previous state. Unintentionally performed hand activations are marked with red ellipses. **c**, Time-series of posterior class probability distribution. Class-specific confidence thresholds are marked with dashed lines.

Overall results for the 12 able-bodied and two amputee participants are reported in Fig. 4. Completion rates and times were used to assess prosthetic control performance. Individual scores are presented on the left-hand side of Fig. 4a and 4b, respectively. Summary scores for the two participant groups are shown on the right-hand side of the graphs. The median completion rates were 95% and 85% for the able-bodied and amputee groups, respectively. Median completion times for successful trials were 37.43 and 44.28 s, respectively. Able-bodied subjects performed on average better than amputees with respect to both metrics, but a formal statistical comparison between the two groups was not possible, due to the small sample size of amputee participants (*n* = 2).

**Fig. 4.**
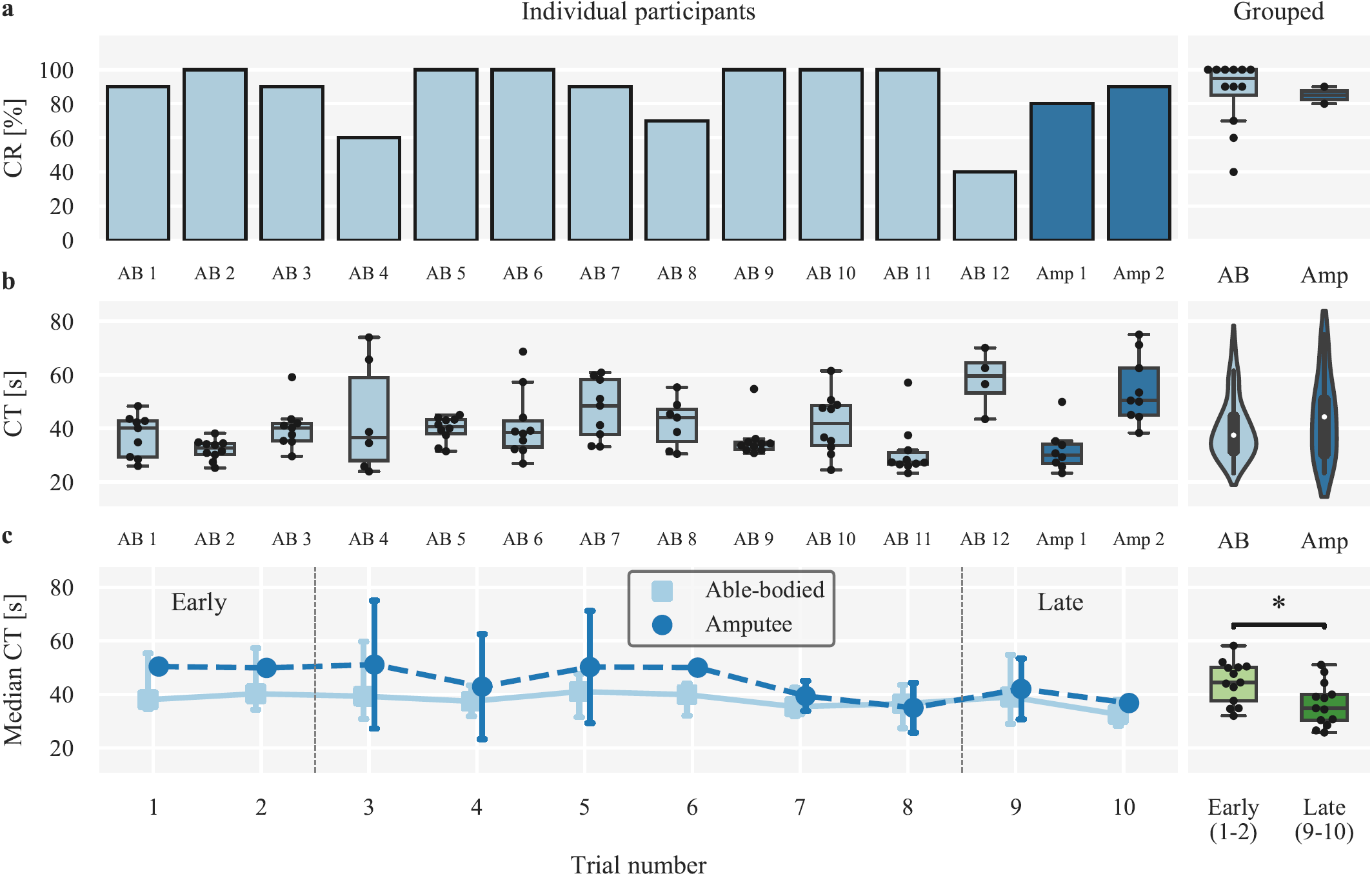
Real-time prosthesis control experiment results. **a**, Completion rates for individual subjects *(left)* and participant groups *(right)*. **b**, Completion times for successful trials. Violin plots, kernel density estimates of sample populations. **c**, Median completion times across participant groups against trial number *(left)* and statistical comparison of early versus late trials *(right)*. Points, medians; error bars, 95% confidence intervals estimated via bootstrapping (1000 iterations); double asterisk, ***p* < 0.05**; CR, completion rate; CT, completion time.

In addition to overall performance, we investigated the effect of task practice on prosthesis control and present the results of this analysis in Fig. 4c. Median completion times are plotted against trial numbers, separately for the two participant groups. We found that median completion times significantly decreased from the first two (*early*) to the last two (*late*) trials (median difference 7.78 s, *p* = 0.007; *n* = 13; Wilcoxon signed-rank test; one able-bodied participant was excluded from this analysis as they did not complete any early trials). The average time elapsed between the initiation of the early and late stages of the experiments (i.e., start of the first and ninth trials, respectively) was 16.08 ± 1.39 min (mean ± s.e.).

The selected sensor pairs for all participants are shown in Fig. 5 using a matrix plot. Red squares indicate selected sensors, whereas blank squares correspond to sensors not available in amputee participants. The last column of the graph shows normalized count for each sensor, computed in the able-bodied population only, as placement locations were different in the amputee participants. Although some sensors were more commonly selected than others (i.e., 2, 6, 8, 10, 15), selection patterns generally varied across individuals.

**Fig. 5.**
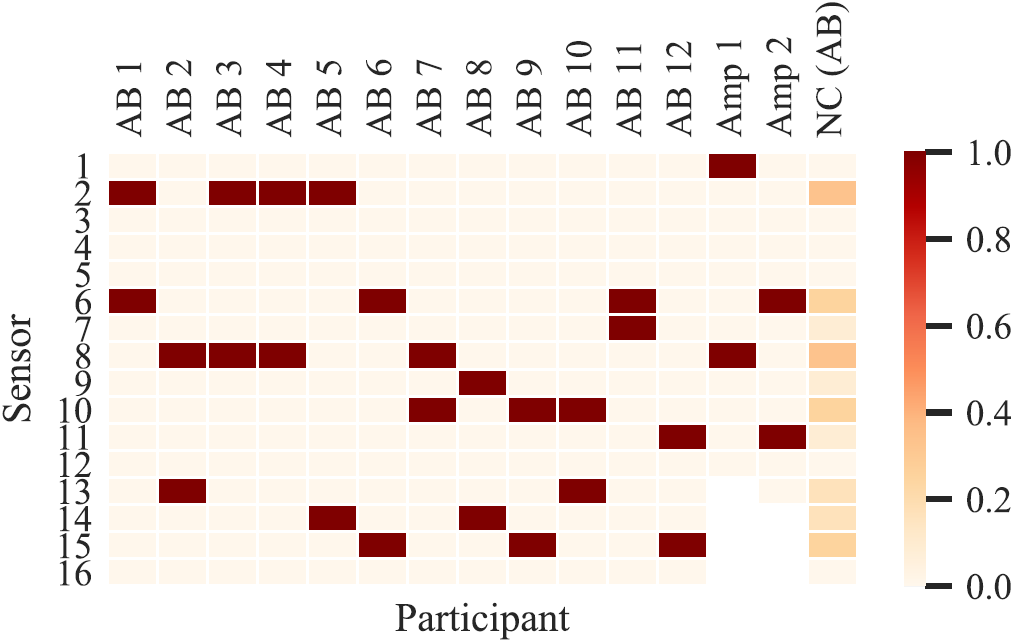
Sensor selection. Red squares indicate the pair of selected sensors for each individual. Blank squares indicate sensors not available in the two amputee participants. The last column shows normalized count for each sensor across the able-bodied population. Sensor locations were shared across able-bodied individuals, but were different for the two amputees. AB, able-bodied; Amp, amputee; NC, normalized count.

An example of confidence threshold selection for one participant is illustrated in Fig. 6 for the “lateral” class. The rejection threshold for the shown example was 0.990 and the corresponding true positive rate was 0.439.

**Fig. 6.**
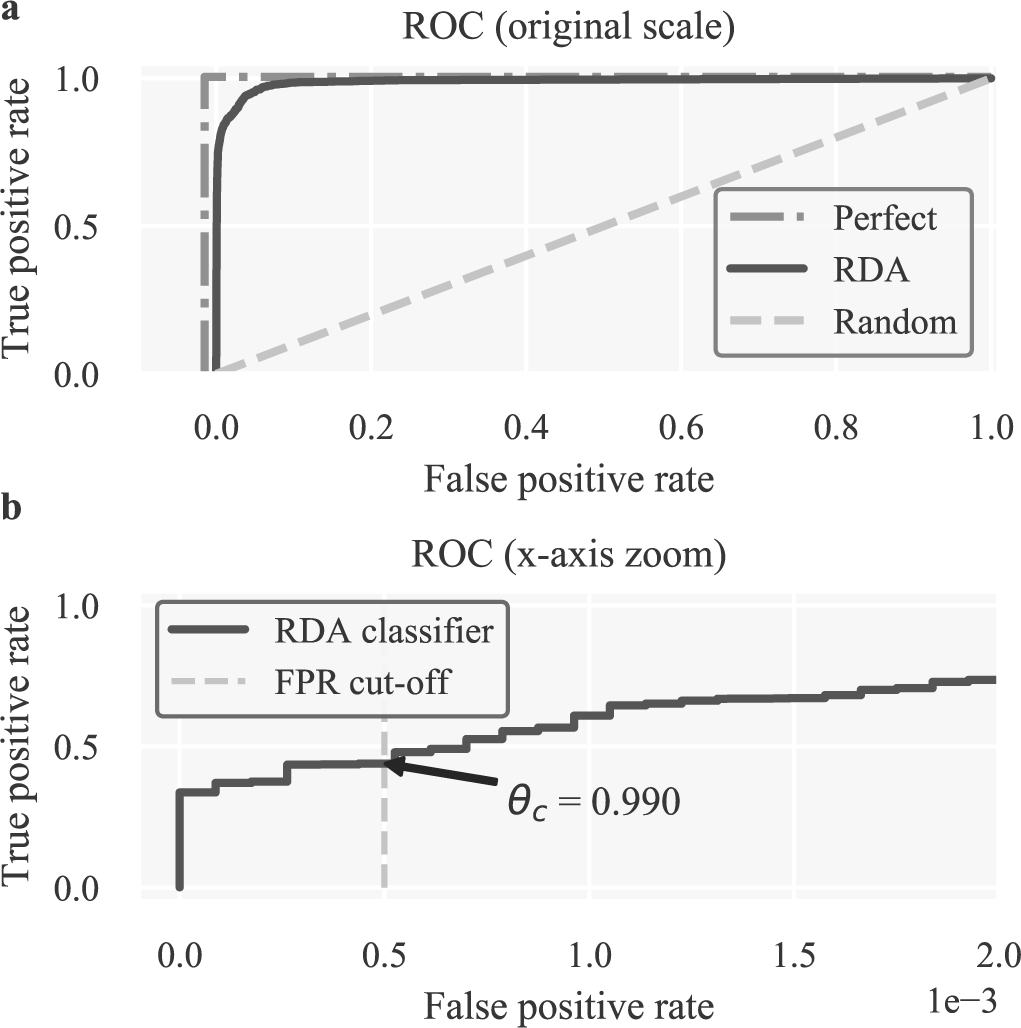
Confidence threshold selection. Class-specific thresholds are selected by transforming the multi-class classification problem into multiple one-vs.-rest problems so that individual ROC curves can be computed. **a**, ROC curves shown for perfect, random, and a representative RDA classifier used in the experiments. **b**, The class-specific confidence thresholds are defined by the points where the respective one-vs.-all ROC curves cross the false positive rate cut-off value of **5 × 10**^**−4**^.

### C. Relationship between offline and real-time performance metrics

We calculated the average balanced offline classification accuracy and cross-entropy loss using 10-fold cross-validation for each subject. The offline scores were then contrasted with the average completion times (across all trials) in the real-time control experiment in a subject-specific fashion. The results of this analysis are reported in Fig. 7 using scatter plots, where each data point corresponds to one individual. Robust linear regression fits using the Huber method (*ϵ* = 1.345, *n* = 14) with 95% confidence intervals are also shown in the same graphs. A positive significant (*p* = 0.04) correlation was observed between cross-entropy loss and completion time. Conversely, the correlation between classification accuracy and completion time was negative and not significant (*p* = 0.10).

**Fig. 7.**
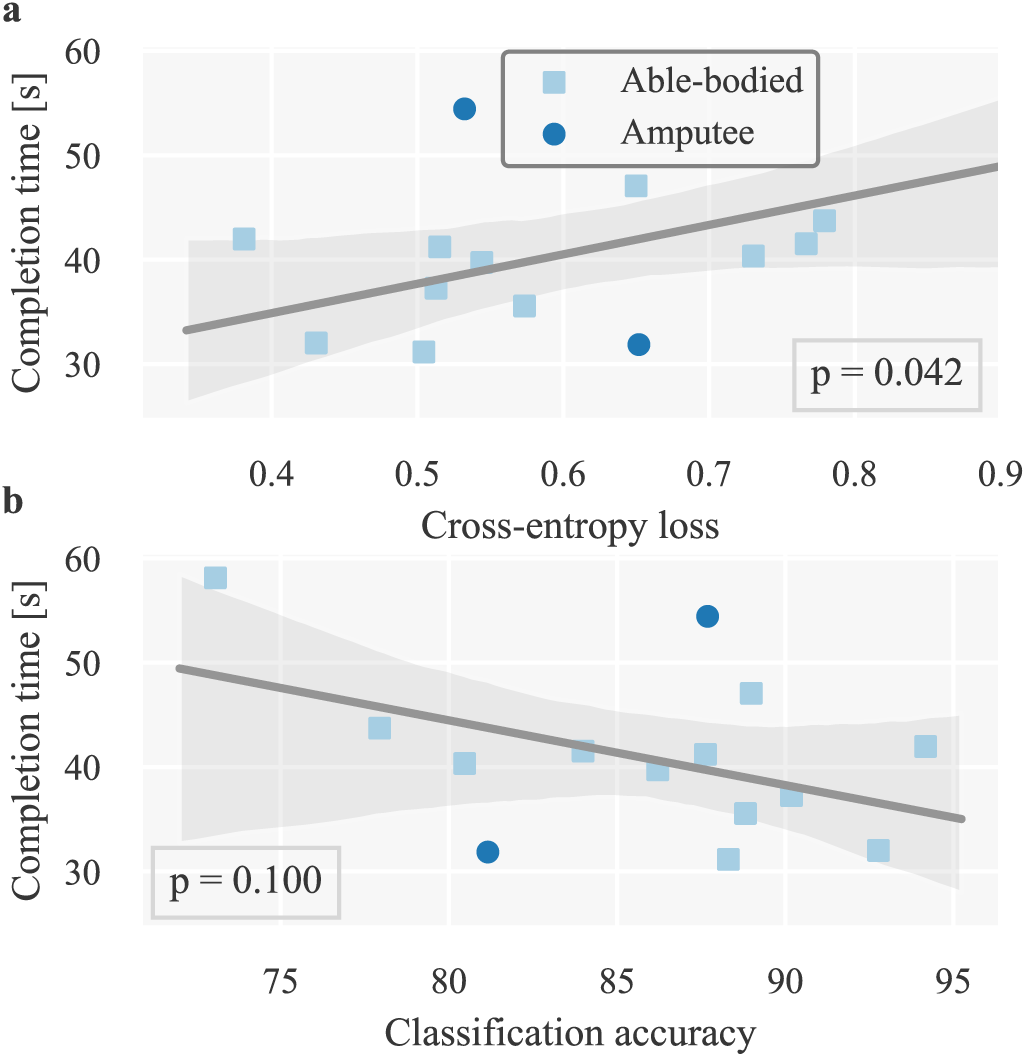
Relationship between offline and real-time control performance metrics. Linear relationships between average completion times and two offline metrics, i.e., **a**, cross-entropy loss and **b**, classification accuracy. Points, individual observations (i.e., participants); lines, robust linear regression fits (Huber method, ***ϵ* = 1.345**); translucent bands, 95% confidence intervals estimated via bootstrapping (1000 iterations).

## IV. DISCUSSION

We have used a real-life object pick-and-place task to demonstrate, for the first time, a proof-of-principle for realtime, classification-based grip control of hand prostheses in transradial amputees using only two EMG-IMU sensors. To tackle this challenging problem, we have proposed a series of novel techniques in terms of sensing, movement intent decoding, and decision making. In the sensing domain, we have made use of our previous finding that the combination of EMG and inertial measurements can improve the performance of classification-based prosthesis control [14]. Although IMUs are typically not available in commercial systems, they are relatively inexpensive components and their integration into existing systems should pose no significant challenges. In fact, some commercial devices already comprise IMUs to monitor prosthesis orientation, but these are embedded in the prosthesis rather than placed on the forearm, as was the case in our study.

For movement intent decoding, we exploited the superior performance of the RDA classifier to LDA [34], especially for a small number of sensors (Fig. 2a, b). This is due to RDA producing quadratic decision boundaries in feature space, while using regularization to avoid the overfitting issues that QDA suffers from. This was verified in the offline analysis, where we observed a decrease in QDA performance as the number of sensors increased; this is a clear sign of overfitting, due to estimating an increasingly larger number of covariance matrix parameters without regularization. By providing a continuum between LDA and QDA, RDA is guaranteed to achieve at least as high performance as that of the two other methods [33].

Notwithstanding the improvement provided by the use of RDA, reducing the number of sensors to only two may inevitably lead to some decrease in classification performance. To pre-empt this potential reduction, we employed a novel confidence-based rejection strategy. We consider this component as paramount for the efficacy of the control scheme proposed here; without it, a substantial number of incorrect classifications might be executed by the prosthetic hand leading to performance deterioration, user frustration, and potentially damage or injury during daily life use [6], [26]. As with previously used algorithms [6], [29], our method was based on discarding predictions that were not made with high confidence using class-specific thresholds [24]. One major difference of our approach was that the function used to estimate the thresholds was based on false positive rate minimization. A quantitative comparison between our approach and previously proposed confidence rejection algorithms was not performed here and is currently seen as a future research direction.

Importantly, the confidence-based rejection step heavily relies on the ability to accurately estimate posterior class probabilities rather than only predicting the most probable class. Taking into consideration that many parameters need to be optimized during training, such as sensor location, classifier hyper-parameters, and rejection thresholds, a metric quantifying performance with respect to the quality of posterior probability estimates is deemed necessary. To that end, we used the cross-entropy loss, in other words, the Kullback-Leibler divergence between the true and estimated multinomial posterior class distributions (see Methods). It is worth noting that the choice of metric directly informs the selection of hyper-parameters; different metrics can often yield utterly different results. By tuning hyper-parameters with respect to minimizing cross-entropy loss, we ensured that the chosen configuration would be optimal with respect to the quality of posterior probability estimates, which plays an important role in ultimate prosthesis control given the final confidence-based rejection stage. Unfortunately, this aspect is often neglected in the field of pattern recognition-based myoelectric control; while most studies involving real-time experiments include some kind of confidence-based rejection, the ability of the decoding algorithm to produce accurate posterior probability estimates is, with very few exceptions [35], not assessed. Instead, the large majority of studies is predominantly concerned with maximizing classification accuracy, despite increasing evidence showing that this metric is not well correlated with real-time control performance [36]–[38]. In our real-time experiments, we found that average completion times were more strongly correlated with cross-entropy loss than with classification accuracy (Fig. 7). It is worth noting, however, that for both metrics, the data points corresponding to the two amputee participants fell outside the respective confidence intervals. We speculate that this discrepancy might be due to different presentation and feedback pathways in individuals with limb loss. However given the small sample size (i.e., number of amputee participants), it is not possible to draw any definitive conclusions. Further investigation with a larger number of transradial amputees may be required to verify whether cross-entropy loss or other probability-related metrics, for example, area under the ROC curve, can provide a reliable estimator of real-time prosthesis control performance.

Moreover, paramount to maximizing decoding performance is tuning the RDA regularization hyper-parameter in a systematic way, for example, using holdout or cross-validation. In addition to the regularization hyper-parameter, sensor selection and rejection thresholds can influence final prosthesis control. Hence, it is crucial to tune them by evaluating performance on a different dataset than the one used for training [39].

For rejection threshold selection using ROC curve analysis, the most commonly used strategies involve either maximizing the vertical distance from a random classifier or minimizing the distance from an ideal classifier [40]. However, neither of the two methods impose a constraint on the resulting false positive rate. This was regarded as a high priority in our case, given the high associated cost of false positive activations which translate into unintended hand activations. To address this issue, thresholds were selected such that the true positive rate was maximized, while constraining false positive rate to be kept lower than a cut-off threshold (not to be confused with the prediction rejection threshold). The value for the false positive rate cut-off threshold was set empirically during pilot trials. An interesting avenue for future research would be to attempt to systematically identify the optimal false positive rate cut-off threshold during real-time myoelectric control. One possible way to achieve this could be by giving the user control over the threshold value, for example via a knob switch, and asking them to select it according to their individual preference. It shall be interesting to investigate whether a shared preference pattern can be observed across different individuals.

Various algorithms have been previously proposed for EMG channel selection, including, but not limited to, exhaustive search (i.e., brute force) [17], [19], sequential forward [18], [23], [41] or backward selection [25], and independent component analysis [21]. We employed the standard sequential forward selection algorithm, mainly because of its speed and efficiency during training. It has been previously demonstrated that despite its simplicity, it can outperform more sophisticated methods, such as the Lasso algorithm [23]. An alternative would have been to optimally select the sensor pair using exhaustive search. In our experience, this approach can only offer a marginal improvement in performance, if any, at the expense of a substantial increase in computational complexity; the number of search iterations scales quadratically in the number of sensors with exhaustive search, as opposed to linearly with sequential selection. Furthermore, it might have been possible to exploit spatial information in EMG and IMU signals to improve performance when using a reduced number of sensors [13]. We did not pursue this aspect here, but we consider it as an interesting avenue for future work.

During offline analysis, we found that the optimal number of sensors for classification may lie in the range of five to seven (Fig. 2a). This is in agreement with previous work investigating EMG channel reduction [14], [18]–[24]. Bearing in mind clinical applications, we sought to investigate whether machine learning-based prosthesis control could be feasible with only two sensors. Contrasting completion times between the current and our earlier work, where we have used an average of four to six sensors in a similar task [14], we note that performance between the two conditions was comparable.

We did not observe any patterns shared across participants in terms of the positions of selected sensors. From a clinical point of view, this finding suggests that a subject-specific approach for sensor position identification may be required. One possible solution to this problem is to adopt the approach considered here; record muscular activity from many sites during an initial screening, and subsequently identify the optimal sensor positions based on a search algorithm. This procedure should however precede socket fabrication, which requires that sensor positions be already established.

It has been previously demonstrated that performance with classification-based controllers can be improved with user practice [17], [42], [43]. In line with previous reports, a significant decrease was observed in completion times between early and late trials (Fig. 4c). The testing phase of the experiment lasted on average 20 minutes, thus it is reasonable to expect that performance could potentially further improve with daily use, provided that the effect of exogenous parameters such as skin condition and sensor position is controlled.

There are a few limitations associated with our study. Firstly, our experimental paradigm was custom, which renders performance comparisons between our approach and other studies somewhat difficult. In the future, it shall be valuable to compare our approach with state-of-the-art algorithms from the literature as well as clinical standards, for example, myoelectric mode switching and body-powered prostheses using standardized clinical tests and metrics. Secondly, although we attempted to simulate a realistic prosthesis control scenario in the lab as closely as possible, wire connections to the hand— for power supply and data transfer—hindered free movement and therefore participants were instructed to remain seated throughout the experiments. A full clinical translation of the proposed paradigm will require an embedded implementation with wireless data transfer and battery-operated components, which will also facilitate the assessment of long-term viability and system performance in a more unconstrained environment.

## V. CONCLUSION

In this study we have provided a pre-clinical proof-of-principle for classification-based multi-grip myoelectric control using only two sensors comprising EMG electrodes and IMUs. Control of a state-of-the-art commercial prosthesis was demonstrated in real-time with able-bodied participants and transradial amputees. The proposed paradigm has the potential to transform existing upper-limb myoelectric systems, subject to minimal modifications, to support intuitive, machine learning-based multi-grip prosthesis control.

## VI. ACKNOWLEDGMENT

The authors thank the two amputee volunteers for participating in this study and Sigrid Dupan for providing comments on an earlier version of the manuscript.

